# Cryo-EM reveals an entangled kinetic trap in the folding pathway of a catalytic RNA

**DOI:** 10.1101/2022.04.05.487152

**Authors:** Steve L. Bonilla, Quentin Vicens, Jeffrey S. Kieft

## Abstract

Functional RNAs fold through complex pathways that can contain misfolded “kinetic traps.” A complete model of RNA folding requires understanding the formation of such misfolded states, but they are difficult to characterize due to their transient and potentially conformationally dynamic nature. We used cryo-electron microscopy (cryo-EM) to visualize a long-lived misfolded state in the folding pathway of the *Tetrahymena thermophila* group I intron, a paradigmatic RNA structure-function model system. The structure revealed how this state forms native-like secondary structure and tertiary contacts but contains two incorrectly crossed strands, consistent with a previous model. This incorrect topology mispositions a critical catalytic domain and cannot be resolved locally, as extensive refolding is required. This work provides a structural framework for interpreting decades of biochemical and functional studies and demonstrates the power of cryo-EM for the exploration of RNA folding pathways.

## INTRODUCTION

To function, RNAs must find their native 3-dimensional (3D) fold among a multitude of alternative conformations in a biologically relevant timescale (*1, 2*). This process is not trivial; the high stability and promiscuity of base-base interactions creates an intrinsic thermodynamic propensity to form non-native contacts, potentially trapping the RNA in stable misfolded states and slowing down the folding process (*1-4*). So-called “kinetic traps” have been detected in the folding pathways of several model RNAs and are more likely in large RNAs with intricate 3D folds (*5-7*). *In vivo*, kinetic traps may be resolved by the action of chaperones that actively misfold the RNA and allow it to refold and/or by the binding of proteins that bias the folding pathway towards the functional fold (*8-10*). Understanding the principles governing formation and resolution of misfolded states is thus a critical step towards a complete model of RNA folding. However, direct visualization of these states has not been generally possible by crystallography because they may be transient and/or conformationally dynamic. Dynamic states may be observed using NMR, but detailed structural information is limited by the size of the RNA. Given the latest advances in the ability of cryo-electron microscopy (cryo-EM) to solve dynamic RNA-only structures (*11-13*), we reasoned that this technique may offer a way to directly observe the 3D structure of RNA folding intermediates, including misfolded states, that have been elusive to structural biology.

The self-splicing *Tetrahymena thermophila* group I intron, the first catalytic RNA discovered, and its multi-turnover ribozyme derivative (herein referred to as TET; Fig. 1A) are well-established model systems used for decades to dissect general principles of RNA folding and catalysis (*14-17*). TET catalyzes the cleavage of a substrate strand using an exogenous guanosine nucleophile (Fig. 1B). This ∼125 kDa ribozyme folds into a compact structure with an internal core of stacked helices stabilized by tertiary contacts between surrounding peripheral domains (Fig. 1C).

**Fig. 1.**
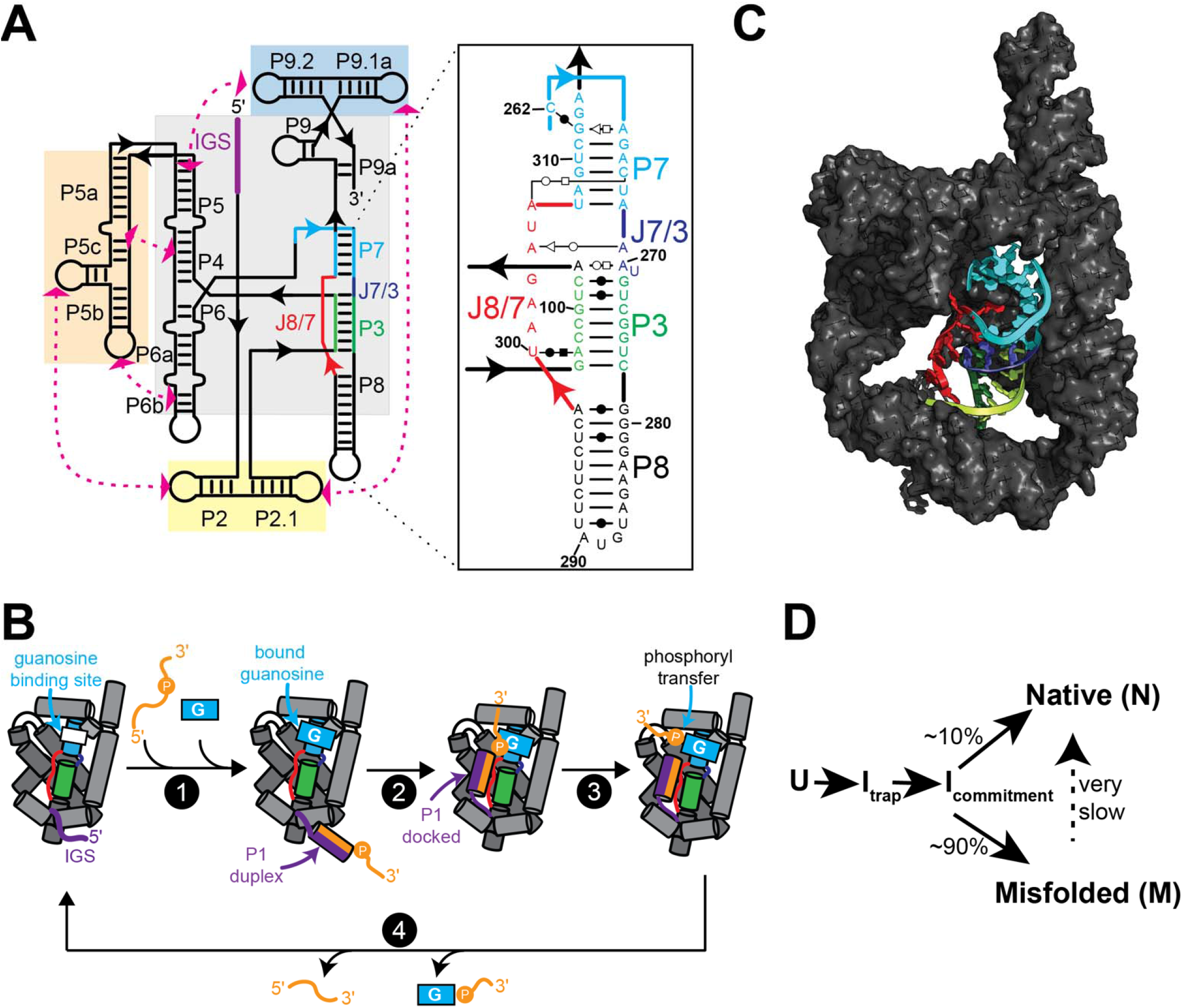
Structure and function of TET ribozyme. (A) Schematic of the secondary structure TET. Paired regions (P) and junctions (J) are named using conventional nomenclature. Grey and colored boxes highlight core and peripheral domains, respectively. IGS: internal guide sequence. Inset shows secondary structure of domains in the proximity of P7. (B) Simplified catalytic cycle of multi-turnover TET ribozyme. For brevity, multiple processes are illustrated as single steps. For a detailed mechanistic description see ref. 15. (1) Oligo substrate anneals to IGS to form P1 and exogenous guanosine binds to pocket in P7. (2) P1 docks into the core. (3) Phosphoryl transfer between guanosine nucleophile and oligo substrate and undocking of P1. (4) Release of products to regenerate *apo* ribozyme. Colors as in (A). (C) Published cryo-EM structure of *apo* L-21 ScaI TET ribozyme in its native folded state (PDB ID: 7ez0). P7, P3, P8, J8/7, and J7/3 are shown in cartoon representation and colored as in (A). (D) Simplified folding pathway of TET ribozyme at 25°C in the presence of 10 mM Mg^2+^. U is the unfolded state. I_trap_ is hypothesized to be a largely structured misfolded intermediate (*21*). I_commitment_ is a common intermediate that precedes commitment to pathways leading to N or M (*21*).

Several intermediates have been identified in the folding pathway of TET, including a long-lived misfolded state (referred to as ‘M’) that can be experimentally accumulated (*18-22*). At standard *in vitro* conditions, ∼10 % of molecules fold directly to the native state (N) and ∼90 % to M (Fig 1D). Refolding from M to N is very slow, in the timescale of hours, suggesting that this process requires considerable structural reorganization (*21, 22*). Consistent with this, solution conditions and mutations that destabilize RNA tertiary structure accelerate re-folding from M to N (*22*). Mutagenesis of the ribozyme core demonstrated that formation of non-native base pairs leads to M; thus, it was initially hypothesized that M contains non-native secondary structure elements near the catalytic center (*21, 23, 24*). Paradoxically, hydroxyl radical and dimethyl sulfate (DMS) footprints of M and N show only minor differences localized mostly to the functionally important P7 helix (Fig. 1A & 1C), suggesting that M and N essentially form the same secondary structure and tertiary contacts (*22*).

To explain the paradoxical nature of M, it was hypothesized that, although M and N form nearly identical structures, two single-stranded elements are crossed incorrectly in M, resulting in a non-native trapped topology that requires extensive unfolding to be resolved (*22*). According to this model, the alternative secondary structure supported by mutations biases the folding pathway toward M and traps the ribozyme in the incorrect topology, but it is later replaced by native-like contacts prior to forming M. Although this hypothesis is consistent with the biochemical and functional data, the 3D structure of the M state has remained elusive, and the topological isomer model remained to be tested directly.

Recently, complete structures of *apo* and *holo* TET ribozymes in the native folded state were solved by cryo-EM (*12*). Concurrent with those studies, we applied cryo-EM to solve the structure of both the N and the M states of TET. By directly observing and comparing these two states, we rationalize decades of studies and paradoxical observations and demonstrate the power of cryo-EM to dissect RNA folding intermediates.

## RESULTS

### Cryo-EM reveals two major conformational states of TET

Previous studies showed that M can be enriched (∼90%) by folding TET at 25°C in the presence of 10 mM Mg^2+^ (*22*). We therefore *in vitro* transcribed and purified the L-21 ScaI ribozyme sequence and folded the RNA using those conditions, then immediately prepared cryo-EM samples and imaged using a 300 kV Krios cryo-EM microscope (Fig. S1; Table S1). *Ab initio* 3D reconstructions and refinements generated maps consistent with the global structure of TET, but the density at the core was not well-defined and suggested a mixture of states. To address this, we performed 3D variability analysis, which uses probabilistic principal component analysis to fit a linear subspace describing variability in the particles (Fig. S2; Movie S1) (*25*). This analysis confirmed that there was conformational heterogeneity in the sample and revealed two conformational classes, with major differences localized to the core of the ribozyme (Fig. 2A). Other volumes generated by particle classification could be assigned to one of the two conformational classes (Fig. S2). The best quality maps from each class were refined to 3.4 Å (N) and 3.9 Å (M) resolution (Fig. 2B & Fig. S4).

**Fig. 2.**
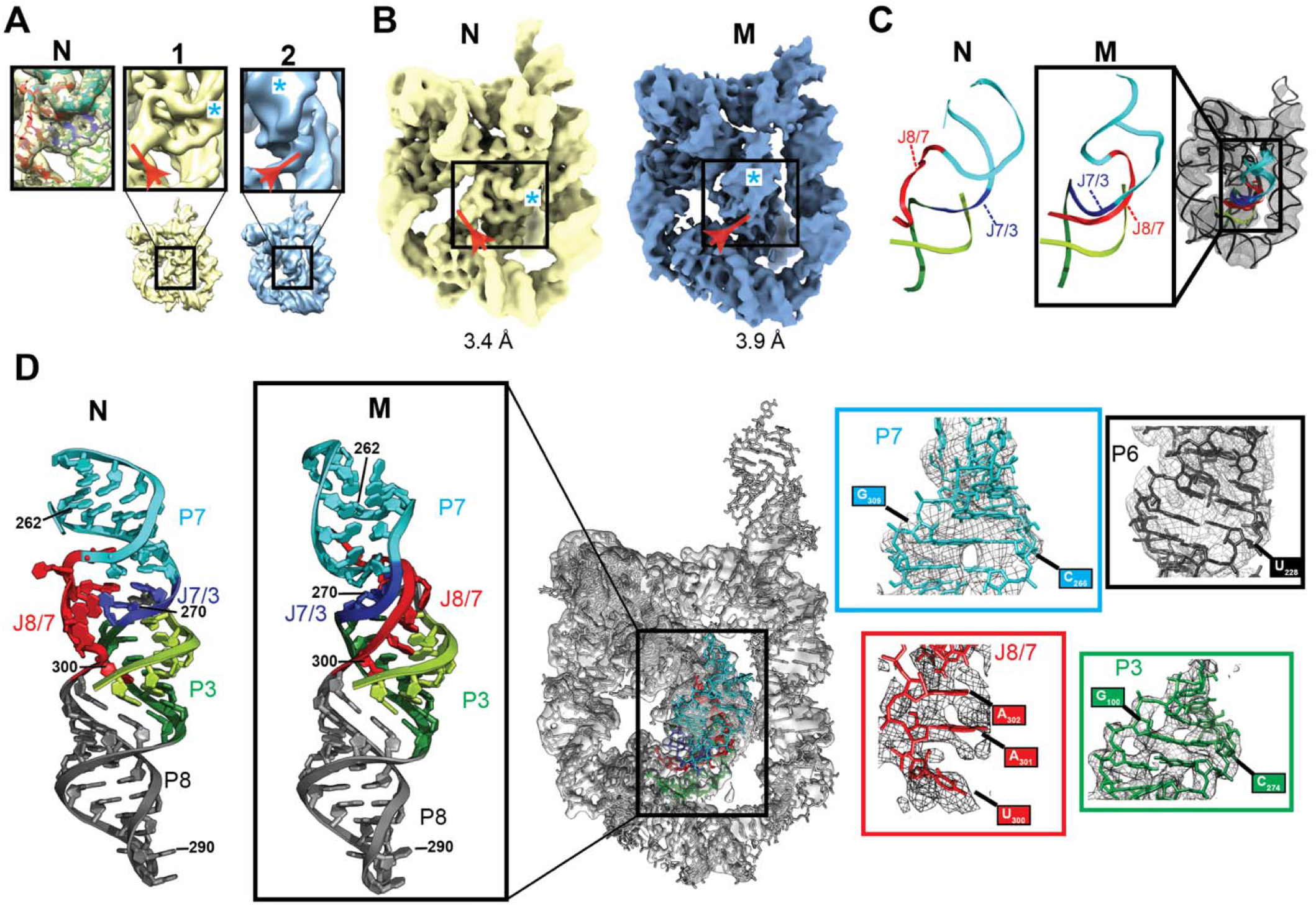
Cryo-EM studies of TET folded at room temperature revealed two distinct conformational states. (A) Particle classification revealed two distinct conformational classes (Maps 1 and 2; Fig. S1 & S2). Major differences within the core are shown in boxes. Density corresponding to J8/7 is marked with a red arrow. Density corresponding to the minor groove of P7 is marked with a cyan asterisk. Left box shows published structure of N fitted to map 1. (B) Maps of N (yellow) and M (blue) states, corresponding to refined versions of maps 1 and 2 in (A), respectively. (C) Models generated by autoDRRAFTER superimposed on cryo-EM map. Representative modelled structure of modelled is enlarged and compared to the same region of N. (D) Atomic model of the full M state after refinements. Refined structure is docked into cryo-EM map (middle). Atomic model of the P7-P3-P8 stack in M is compared to that of N (far left) to highlight structural differences between these two states. Close-ups show examples of structural models fitted to density (right panels).

The first class (Fig. 2A & 2B, yellow) matches the published cryo-EM map of TET in the N state (correlation: 0.94) and the published structure docks well into the map without additional refinements (avg. CC_mask_: 0.77; Fig. S3). The map-model correlation per residue was essentially the same with the published map and with our map (Fig. S3). Therefore, this class represents a population of N in the sample, consistent with previous studies showing that a fraction of molecules fold directly to N under the experimental conditions (Fig. 1D) (*21, 22*). In contrast, the second class (Fig. 2A & 2B, blue) does not fit the structure of N and revealed major differences in the core near the functionally important P7 helix, which contains the conserved guanosine-binding site, indicating that this class likely corresponds to the M state.

### 3D structures reveal topological differences between the N and M states

The core of TET contains two sets of stacked helices: P4-P5-P6 and P3-P7-P8-P9 (Fig. 1A; grey box). J8/7 is an unpaired stretch of seven nucleotides that links P7 to P8 and makes contacts with the P3 helix (Fig. 1A). Comparison of N and M maps suggested that the peripheral domains and the P4-P5-P6 stack of the core were similar between the two conformations, although with minor differences discussed below. In contrast, the maps suggested that J8/7 extends in a different direction in each map (Fig. 2A & 2B, red arrow) and that P7 is rotated in M relative to its position in N (Fig. 2A & 2B, cyan asterisk). To learn more about these conformational differences, we fitted an atomic model into the density of M. As the resolution was not sufficiently high for manual fitting, we used autoDRRAFTER (*13*) for map-guided computational modelling of the core of M, keeping the peripheral elements, the P4-P5-P6 stack, and the secondary structure fixed during modelling (Fig. 2C & Fig. S5). The full structure of M was refined using Phenix and COOT modelling software (Fig. 2D and Fig. S5) (*26, 27*). The final structure fits well into the map (CC_mask_: 0.74) without major steric clashes (clash score: 6.61). While the moderate resolution of the map in the core does not allow atomic-level precision, the strand topology and global structure of M and its comparison to the known structure of N are clear and provide important insights.

The final model of M reveals a core topology that diverges from that of N (Fig. 2C & 2D). Whereas in N the single strand J8/7 is “on top” of J7/3 (which joins helices P7 and P3) as seen from the orientation shown in Fig. 2C, in M J8/7 passes “under” J7/3. This entanglement of J8/7 is accompanied by a ∼90° rotation of P7, likely induced by the new position of J8/7 that constrains proper placement of P7 (Fig. 2D). In contrast, helix P3 is essentially unmoved relative to its native position (Fig. 2D). Consistent with previous observations, long-range tertiary contacts are maintained, even though large rearrangements occur within the core. Remarkably, these observations are consistent with the topology isomer hypothesis discussed above (*22, 28*), although the specific topological differences and the identities of the entangled strands differed from those predicted. Visual inspection shows that this topological error cannot be resolved locally and requires major unfolding of the ribozyme (*vide infra*), consistent with the hours-long time scale at which the M to N transition occurs.

The structure of M is consistent with previous studies showing a strong link between the formation of an alternative P3 (alt-P3) secondary structure and formation of M (*23, 24*). In alt-P3, nts 303-306 in J8/7 are proposed to base pair with nts 271-274, effectively preventing formation of the long-rage native P3 (Fig 1A). Based on the nearly identical chemical probing footprints of M and N, it was later proposed that formation of alt-P3 traps the ribozyme in the incorrect topology, but alt-P3 is later replaced by the native P3 (*22*). The map of M, although not allowing unambiguous positioning of every nucleotide, supports this model. In M, J8/7 docks into the major groove of P3 and is near nts 271-274 where it could readily pair to form alt-P3 (Fig. 3A). Thus, only local rearrangements are required to transition from an intermediate state with alt-P3 to the M state. Although we cannot make any conclusions about the conformation of these elements before M is formed, the structure suggests how formation of alt-P3 could position J8/7 in its final trapped position in M.

**Fig. 3.**
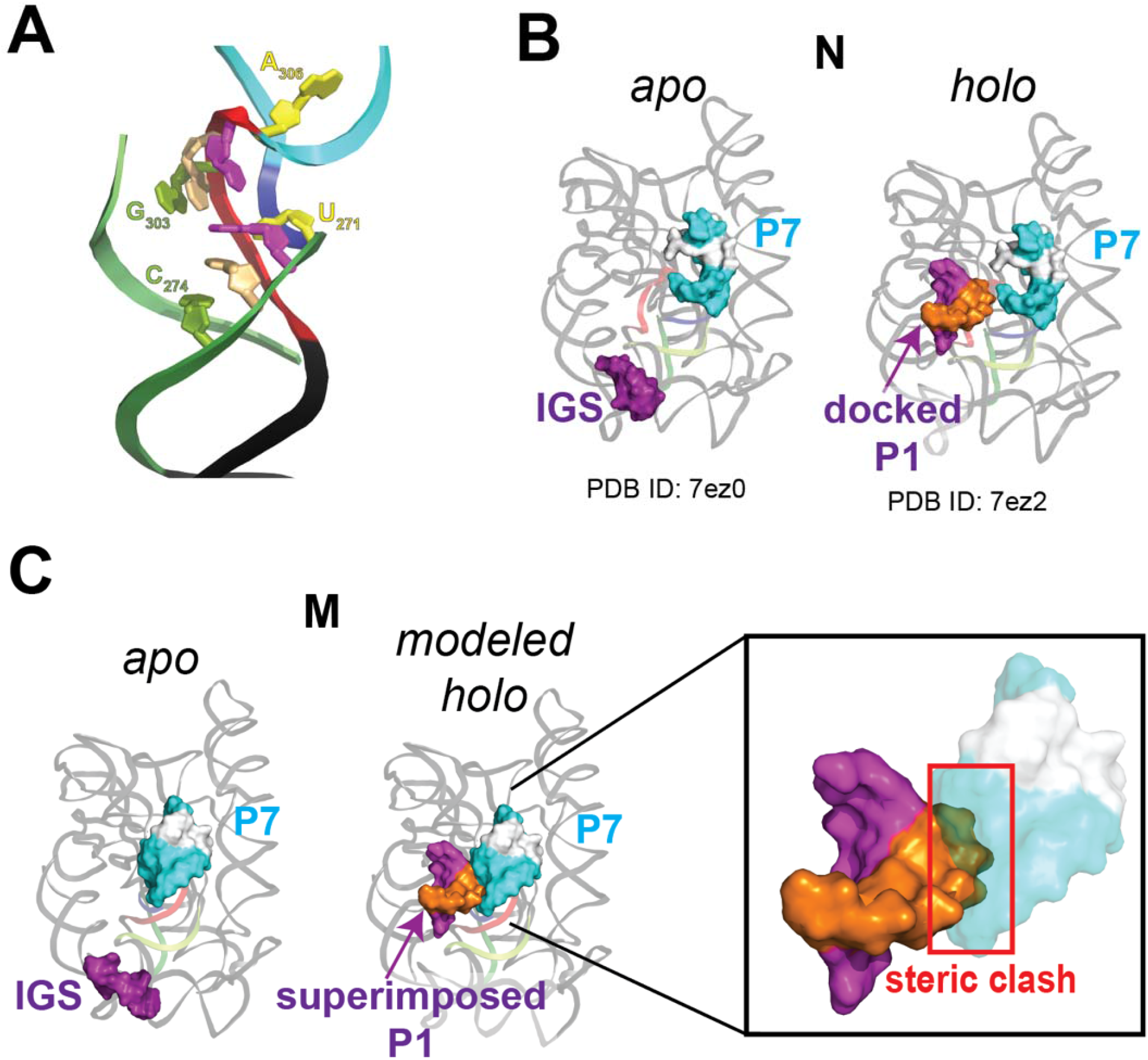
Structure of M provides insights into ribozyme misfolding and catalysis. (A) Nucleotides predicted to base pair in alt-P3, leading to the formation of M. Backbone is colored as in Fig. 2. Bases predicted to pair in alt-P3 are the same color. (B) Published cryo-EM structures of *apo* L-21 ScaI (PDB ID: 7ez0) and *holo* L-16 ScaI (PDB ID: 7ez2) TET ribozymes folded in the native functional state. For clarity, helix P10 and additional base pairs formed by a second oligo substrate in the published structure (*12*) are not shown. P7 and IGS are rendered as surfaces and nucleotides in P7 that form the binding pocket for guanosine are colored white. (C) Superposition of docked P1 (extracted from PDB ID 7ez2) on structure of M state. Inset shows steric clash predicted to block docking of P1 into the core of the M state.

### The M state structure explains its decreased catalytic activity

Given the similarity of the structures of N and M suggested by chemical probing, the difference in their catalytic activity was puzzling, but the 3D structure of M now resolves this paradox. As shown in a simplified scheme of the TET catalytic cycle (Fig. 1B), the P1 duplex – formed by base pairing of the substrate strand to the internal guide sequence (IGS) – docks into the core, positioning the 5’ splice site in proximity to the bound guanosine nucleophile. Structures of N with and without bound substrates showed that the core is globally preorganized for P1 docking (Fig. 3B) (*12*). This is supported by solution small angle X-ray scattering (SAXS) data, which are consistent with no major changes in the overall structure of the ribozyme upon substrate binding (*29*).

In M, the P1 docking site is partially occupied by P7, which is rotated ∼90° and moved by several Å relative to N (Fig. 3C). Thus, the rotated P7 effectively blocks P1 docking, which is an essential step early in the catalytic cycle. Although particle variability analysis (Movie S1) and the lower local resolution of P7 (Fig. S7) suggest that P7 is conformationally dynamic, this helix is unable to rotate to its native position without major rearrangements due to constraints imposed by the connecting strands. Remarkably, it was previously hypothesized that the inactivity of M was caused by rearrangements preventing P1 docking, based on a cleavage assay of a 3’-splice site mimic oligo that does not require P1 docking (*22*). The 3D structure of M strongly supports this hypothesis. Moreover, in the context of the complete group I intron, and based on a structure of TET mimicking the second step of splicing (*12*), the position of P7 would also interfere with the formation of P10 prior to the second transesterification reaction, making M incompetent for self-splicing. Finally, as the guanosine-binding site is contained within P7, the large rotation and displacement of this helix would cause the bound guanosine to be in a different position relative to N, even if P1 was able to dock. Overall, the alternative position of P7, which is coupled to the entangled J8/7, explains the functional differences between N and M.

### M contains small differences in the relative positions of the peripheral domains

Although large differences between M and N are observed at the core, their peripheral domains are very similar (RMSD_backbone_: 2.43 Å; Fig S6). Importantly, M forms native long-range tertiary contacts consistent with previous functional studies of ribozyme mutants (*22*). However, there are small differences in the relative position of the peripheral domains, presumably arising from the distinct core conformations. To better visualize these differences, we superimposed the P4-P6 domains (nts. 107 to 258) of N and M, which are nearly identical in the two states (RMSD: 0.995 Å; Fig S6). With the P4-P6 domain superimposed, the backbone of other peripheral domains is misaligned by up to ∼9 Å, most noticeably in P2 and P9 (Fig. S6). Interestingly, previous SAXS studies suggested a 10% increase in the radius of gyration (R_g_) of M versus N (*29*). We do not observe significant differences in compaction and both structures have essentially the same R_g_ of ∼38 Å. However, the local resolution distributions are very different between N and M, suggesting differences in global flexibility (Fig. S7). In particular P9, which is conformationally dynamic (*12*), appears to be much more flexible in M relative to N, as inferred by the local resolution of this domain (Fig. S7). Thus, differences in R_g_ observed by SAXS may be reflective of differences in the direction and/or magnitude of these dynamics, the presence of other less-compact conformational species in the SAXS sample, or of limitations in the accuracy of R_g_ estimates from SAXS profiles.

### A model for disentangling the topological error

Resolution of the topological error in M cannot occur with only local rearrangements, raising the question of how much unfolding is needed to allow the transition from M to N. Although our data cannot establish the mechanism of this transition, comparing the structures of M and N in the context of previous functional and biochemical studies can provide insights. We generated a model for the transition of M to N that requires minimal secondary structure disruption and that is consistent with previous studies (Movie S2). In this model, tertiary contacts between peripheral domains and the long-range base pairs that form P3 break to allow rotation of a long hairpin that includes P7, J8/7, J7/3, nts 272-278 of P3, and P8 (Movie S2). P7 rotates ∼90° to reach its position in the native state, while P8 undergoes a ∼360° rotation that is allowed by the flexibility of single strands J8/7, J7/3, and nts 272-278. After these rotations, refolding P3 and docking of the tertiary contacts results in formation of N (Movie S2). This model shows that there are paths from M to N that do not require extensive breaking of secondary structure. This model is also in agreement with observations that destabilizing tertiary interactions between peripheral domains and/or base pairs in P3 increase the rate constant for the transition from M to N (*22, 28*). Further, because minimal disruption of secondary structure is required, the model is consistent with the ability of M to convert to N at physiological temperatures, albeit slowly.

## DISCUSSION

RNA folding pathways are complex, with multiple branching points and intermediates. For decades, researchers have applied a wide array of diverse tools to detect folding intermediates and misfolded states in the folding of model RNAs. These techniques include quantitative thermodynamic and kinetic analysis, single-molecule imaging, X-ray scattering techniques, and electrophoretic mobility assays, among others (*30*). Pathways deduced by these techniques have proven invaluable to our understanding of RNA folding and can generate quantitative, testable predictions. What has been lacking is a 3D view of the conformational species populating the pathways. Such information would provide a structural framework for interpreting decades of functional and biochemical studies and generate additional hypotheses, ultimately enhancing our predictive understanding of the folding process. The scarcity of 3D structural information is in part due to a lack of tools to readily explore the structures of these perhaps transient and/or conformationally dynamic states. Building on previous functional and biochemical studies and taking advantage of the recent advances in cryo-EM, we used the *T. thermophila* group I intron ribozyme as a model for RNA misfolding and solved the structure of a long-lived misfolded intermediate state that had remained mysterious for decades. In so doing, we demonstrate the power of cryo-EM as a tool to explore dynamic folding pathways of complex, functional RNAs.

The term ‘misfolded RNA’ might suggest the formation of non-native secondary structures or global changes in tertiary structure. While such states can form, in the case of the long-lived misfolded state of the *T. thermophila* group I intron, referred to as the ‘M’ state, misfolding is generated by a pair of incorrectly crossed single strands that cause a topological error within a fold with native-like secondary structure and tertiary contacts. This topological error results in the large rotation of a functionally important domain, effectively preventing organization of the catalytic site and rendering the M state functionally incompetent. The question of how often topological errors arise within the folding pathways of other complex RNA folds is not clear, but the structure of the M state model provides an example that will facilitate future explorations.

Given the inherent thermodynamic propensity of RNAs to misfold, it is remarkable that biology has found ways to produce functional RNAs in time scales that are consistent with life. In part, kinetic traps may be alleviated *in vivo* by the work of chaperones with helicase activity and/or the stepwise binding of proteins that guide the folding process. Additionally, co-transcriptional folding may bias the folding pathway. A mechanistic understanding of these complex processes requires the understanding of the inherent folding properties of RNA in isolation. Experimental and computational advances have turned cryo-EM into a premier tool to solve the high-resolution native structures of biological macromolecules. Ongoing developments in time-resolved cryo-EM promise dissection of transient intermediate structures (*31*). Here, we demonstrate the application of conventional cryo-EM to understand the structure of a transient but long-lived misfolded intermediate in a complex, folded RNA. This state was not directly observed for many years, in part due to limitations of other structural methods. Now, direct observation of the state provides strong evidence in support of previous models proposed based on functional and biochemical data (*22, 28*). The remarkable fact that these models were largely correct, even in the absence of a 3D structure, illustrates the power of rigorous quantitative biochemistry and biophysics to sort out complex problems in RNA structure and folding, and the natural marriage between these methods and cryo-EM.

## Data availability

Structures and cryo-EM maps will be deposited in online repositories before publication.

## ACKNOWLEDGEMENTS

Theo Humphreys (PNCC) and Eduardo Romero (UC Anschutz) assisted with microscope operation. Dave Farrell provided support of computational resources The authors thank members of the Kieft Lab, Raghuvir Sengupta, and Dan Herschlag for helpful discussions and David Constantino for proofreading. This work was funded by National Institutes of Health grant R35GM118070 to J.S.K. and by a HHMI Hanna H. Gray fellowship awarded to S.L.B. A portion of this research was supported by NIH grant U24GM129547 and performed at the PNCC at OHSU and accessed through EMSL (grid.436923.9), a DOE Office of Science User Facility sponsored by the Office of Biological and Environmental Research.

## MATERIALS AND METHODS

### Preparation of folded TET RNA

We ordered a DNA fragment (gBlocks; Integrated DNA Technologies) containing the L-21 ScaI TET sequence flanked by the T7 RNA polymerase promoter sequence. The DNA fragment was PCR amplified, purified using the GeneJET PCR Amplification Kit (Thermo Scientific), and used as a template for *in vitro* transcription.

TET RNA was transcribed in a 250 µl reaction containing 6 mM of each NTP (ATP, CTP, GTP and UTP), 60 mM MgCl_2_, 30 mM Tris pH 8.0, 10 mM DTT, 0.1% spermidine, 0.1% Triton X-100, 0.24 U/µl RNasin RNase inhibitor (Promega), ∼ 0.14 mg/ml T7 RNA polymerase (in-house prepared), and 37 µM DNA template. The reaction was incubated at 37°C for 3 hours and ethanol precipitated overnight. RNA was purified by denaturing gel electrophoresis (5% polyacrylamide) and by AMPure XP magnetic beads (Beckman Coulter). The purified RNA was buffer exchanged three times into RNase-free water using 30 kDa cutoff Amicon centrifugal filters (Millipore). The RNA was stored at -20°C until preparation of cryo-EM grids.

Prior to preparation of cryo-EM grids, the RNA was folded in 10 mM MgCl_2_ and 50 mM NaMOPS (∼30 mM Na^+^) pH 7.0 by incubating at 25°C for 15 minutes. These conditions accumulate the misfolded state of the ribozyme (*22*).

### Cryo-EM grid preparation

A Gatan Solarus Model 950 advanced plasma system was used to clean C-Flat holey carbon grids (hole size: 1.2 µm; spacing: 1.3 µm; mesh: 400; Electron Microscopy Sciences). Settings used for the plasma cleaning were ‘Cleaning time’: 6 seconds; ‘Vacuum target’: 70 mTorr; ‘Vacuum range’: 0 mTorr; ‘Pumping switch point’: 20 Torr; ‘Turbo pump speed’: 750 Hz; ‘O_2_ gas flow’: 27.5 sccm; ‘H_2_ gas flow’: 6.4 sccm; ‘Air gas flow’: 0.0 sccm; ‘Gas flow timeout’: 20 seconds; ‘Forward RF target’: 50 W; ‘Forward RF range’: 5 W; ‘Maximum reflected RF’: 5 W; ‘RF tuning timeout’: 4 seconds; ‘RF tuning attempts’: 3. Then, 3 µl of folded RNA solution (final concentrations: 16.8 µM RNA, 10 mM MgCl_2_, and 50 mM NaMOPS (∼30 mM Na^+^) pH 7.0) was deposited on each grid and a FEI Vitrobot Mark IV was used to plunge-freeze the grids using liquid ethane. The humidity of the Vitrobot chamber was set to 100% and the temperature to 4°C; blot force was set to –5; blot time was 2.5 seconds with 0 seconds wait time. Filter paper was used for blotting (Prod No. 47000-100, Ted Pella, INC.).

### Data collection and analysis

Data were collected at the Pacific Northwest Cryo-EM Center (PNCC) with a 300 kV Thermo Scientific Krios TEM in super resolution mode (pixel size: 0.5395 Å), equipped with a Falcon 3 direct electron detector (DED) and a Bioquantum K3 imaging filter, and using SerialEM data collection software. We collected 6222 movies (46 frames) with a total dose of 32 electrons/Å^2^ and a defocus range of 0.8 to 2.0 µm (Table S1).

Data were processed using cryoSPARC (Fig. S1 & S2) (*32*). Imported movies were subjected to patch motion correction and CTF estimation with default parameters (Fig. S1). Micrographs were curated to eliminate those with damaged areas, excessive ice contamination, and/or poor CTF estimation, resulting in 5709 curated micrographs. Automated particle picking and extraction (extraction box size: 480 pix; Fourier crop to box size: 256 pix) resulted in 4,272,705 putative particles. To remove junk, three rounds of 2D classification were performed (number of classes: 200; circular mask diameter: 170 Å; final full iterations: 2; online-EM iterations: 40; batchsize per class: 500). In the last two rounds, the initial classification uncertainty factor was set to 10 to obtain a higher diversity of good classes. A total of 803,573 particles remained after 2D classification.

The particles were used to build three *ab initio* models. Two of the models (88% of particles; Fig. S1) displayed RNA features and global structures consistent with TET. The third map (12% of the particles; Fig. S1) was used as a “sink” for removal of junk and suboptimal particles in three rounds of 3D classification (heterogeneous refinement), as done previously (*11*). A total of 650,948 particles remained after 3D classification (n_1_ = 285,646; n_2_ = 365,302; Fig. S2A). The two TET maps were refined using ‘homogeneous refinement’ with default parameters. To resolve remaining conformational heterogeneity, the refined particles were subjected to 3D particle variability analysis (number of modes to solve: 2; filter resolution: 6 Å) (*25*) to generate four “frames” along the first principal component for each of the two original maps (Fig. S2A). Total particles (n = 650,948) were re-classified between the eight frames and refined (volumes 1-8; Fig. S2A). The maps were inspected and classified into two major classes, based on differences in core density (Fig. S2B). Maps with the highest resolutions (volumes 5 and 7; Fig. S2A) were subjected to an additional (non-uniform) refinement. The final maps of N and M were refined to 3.4 Å (n_N_ = 92,828 particles) and 3.9 Å (n_M_ = 98,071 particles), respectively.

### Structural modeling of M state

A schematic of the methods used for structural modelling is provided in Fig. S5. The structure of the L-21 ScaI TET ribozyme (PDB ID: 7ez0), with Mg^2+^ ions removed, was docked into the 3.9 Å resolution map of the M state, using UCSF Chimera (Fig. S5A) (*33*). The docked structure was then imported into Phenix (*26*) and a real-space refinement was performed (macro cycles: 1), constraining the secondary structure. Nucleotides corresponding to P7, P3, J8/7, J7/3 and connecting junctions were removed from the structure and were modeled using ROSETTA RNA fragment assembly implemented in autoDRRAFTER (Fig. S5B) (*13*). The truncated structure (containing the peripheral domains and P4-P6, which fit well into the map) was kept constant during modelling and was provided as an input to autoDRRAFTER, along with the sequence and secondary structure of TET and the 3.9 Å map. The protocol described under ‘Manually setting up an autoDRRAFTER run’ in the ROSIE web server was followed (*34*). Three rounds of autoDRRAFTER modeling were performed (cycles: 30000; number of models: 2000). The modelled structures converged towards a common topology (Fig. S5A). The top ten models were inspected in the context of the map; all ten models fit equally well and one of them was chosen for further refinements. The model was imported into Phenix for a real-space refinement (macro cycles: 5) with secondary structure constraints. Coot (*27*) was used to make minor adjustments to the structure around J8/7 and a final real-space refinement was performed in Phenix (macro cycles: 5). Structures were visualized using Pymol (Schrödinger, Inc) and UCSF ChimeraX (*35*). Radius of gyration was calculated using Crysol (*36*). To generate the model for transition from M to N (Movie S2), we manually modified structures using Pymol to generate intermediate structures. The geometries of the intermediate structures were regularized using Phenix geometry minimization tools and molecular dynamics flexible fitting in Namdinator (*37*), using synthetic electron density maps generated in Chimera as inputs. The intermediate structures were interpolated using ‘morph conformations’ (interpolation method: corkscrew; interpolation rate: linear) in Chimera to generate Movie S2.

## Supplementary Materials for

### Other Supplementary Materials for this manuscript include

Movie S1: Conformational variability analysis

Movie S2: Possible M to N refolding pathway.

**Fig. S1.**
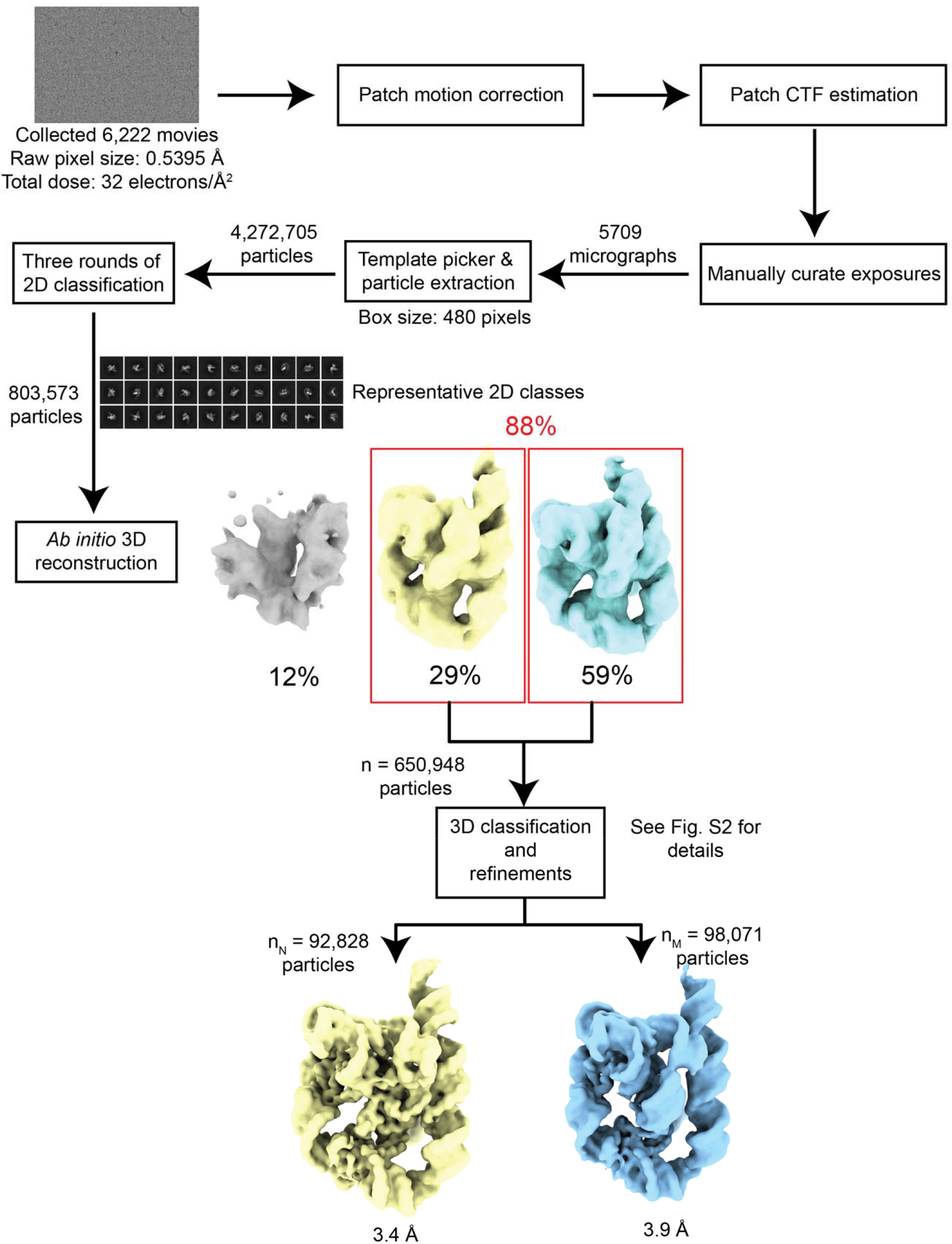
Data analysis workflow. Data were collected with a 300 kV Thermo Scientific Krios TEM in super resolution mode and processed with cryoSPARC (*32*). Parameters are described in Materials and Methods. As done previously (*11*), several rounds of 3D classification were performed to increase the quality of the maps, using a bad *ab initio* reconstruction (grey volume) as a “sink” for suboptimal particles (see Materials and Methods). Details of 3D classifications and final refinements are provided in Fig. S2.

**Fig. S2.**
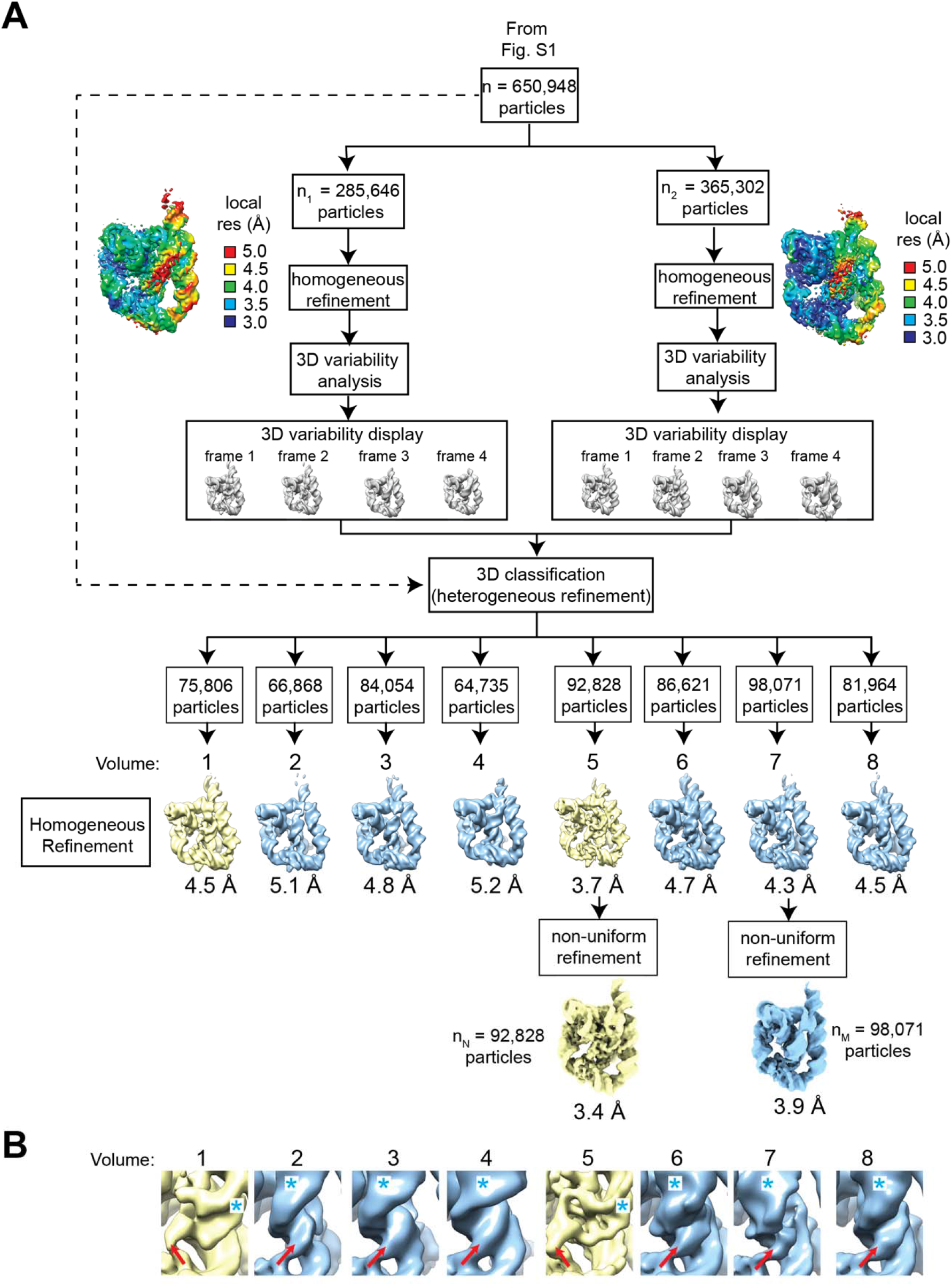
Two ribozyme core conformations revealed by particle variability analysis. (A) cryoSPARC 3D variability analysis (*25*) was used to resolve conformational heterogeneity. Two *ab initio* models consistent with TET structure (Fig. S1) were refined and used as inputs for variability analyses. Four frames along the first principal component were generated for each input map. The total particles (n = 650,948) were re-classified across the eight frames. (B) Close-up of the cores of refined maps generated variability analyses. Inspection of the maps revealed two conformational classes (yellow vs. blue maps). Major differences were observed in the direction of J8/7 density (red arrow) and orientation of the P7 minor groove (cyan asterisk).

**Fig. S3.**
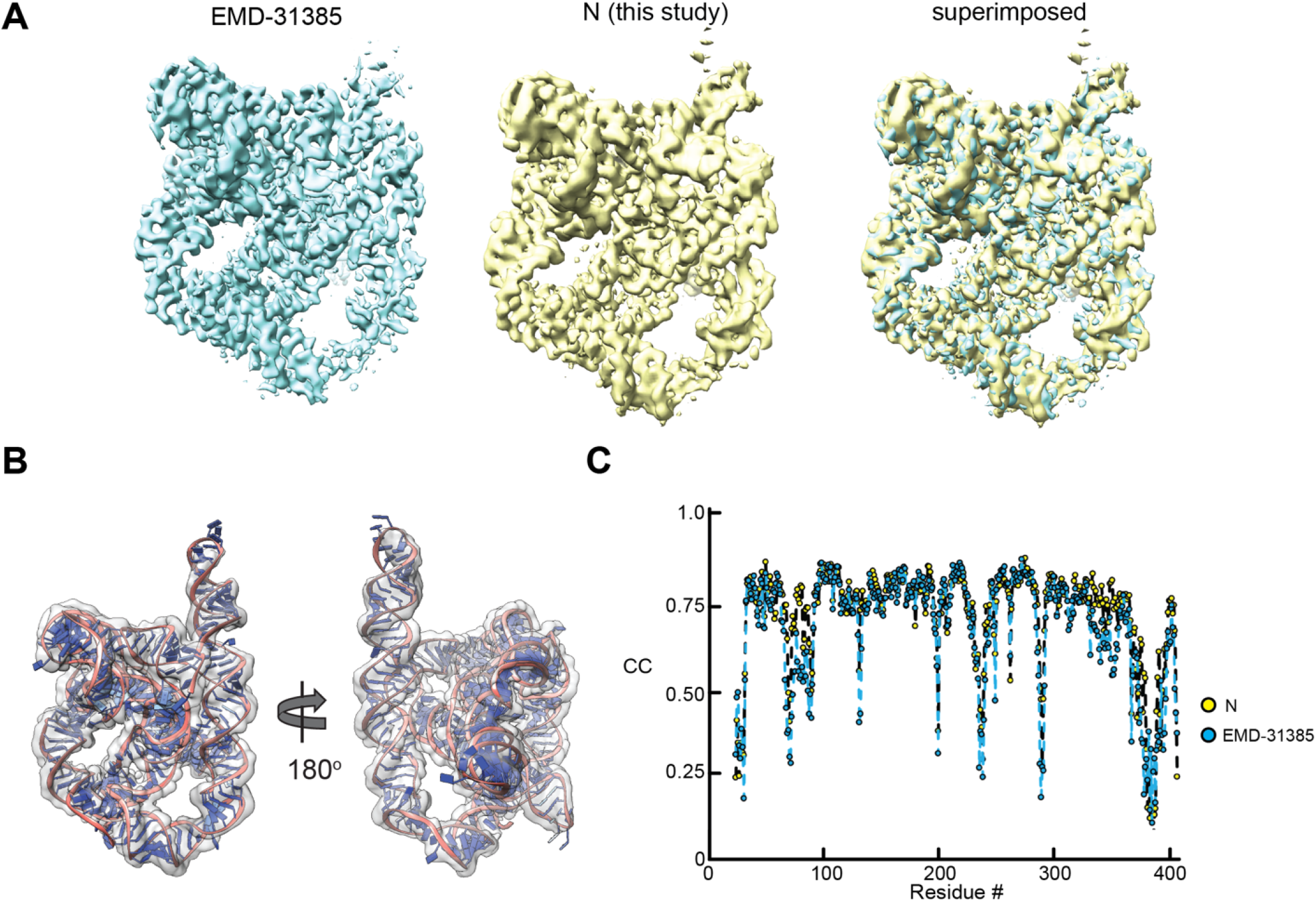
Map of N agrees with published structure of *apo* L-21 ScaI TET ribozyme. (A) Comparison of one of the conformations revealed in this study (folded at 25°C) to one generated in previous studies (folded at 50°C) (EMD-31385) (*12*). The correlation between the maps is 0.94, as calculated with UCSF Chimera (*33*). (B) Published structure of TET in N state (PDB ID: 7ez0) docked into map of N generated in this study (CC_mask_ = 0.77). (C) Correlation per residue was similar between the published map of N (EMD-31385, blue) and the published structure (PDB ID: 7ez0) and the map of N generated in this study (yellow) and the published structure.

**Fig. S4.**
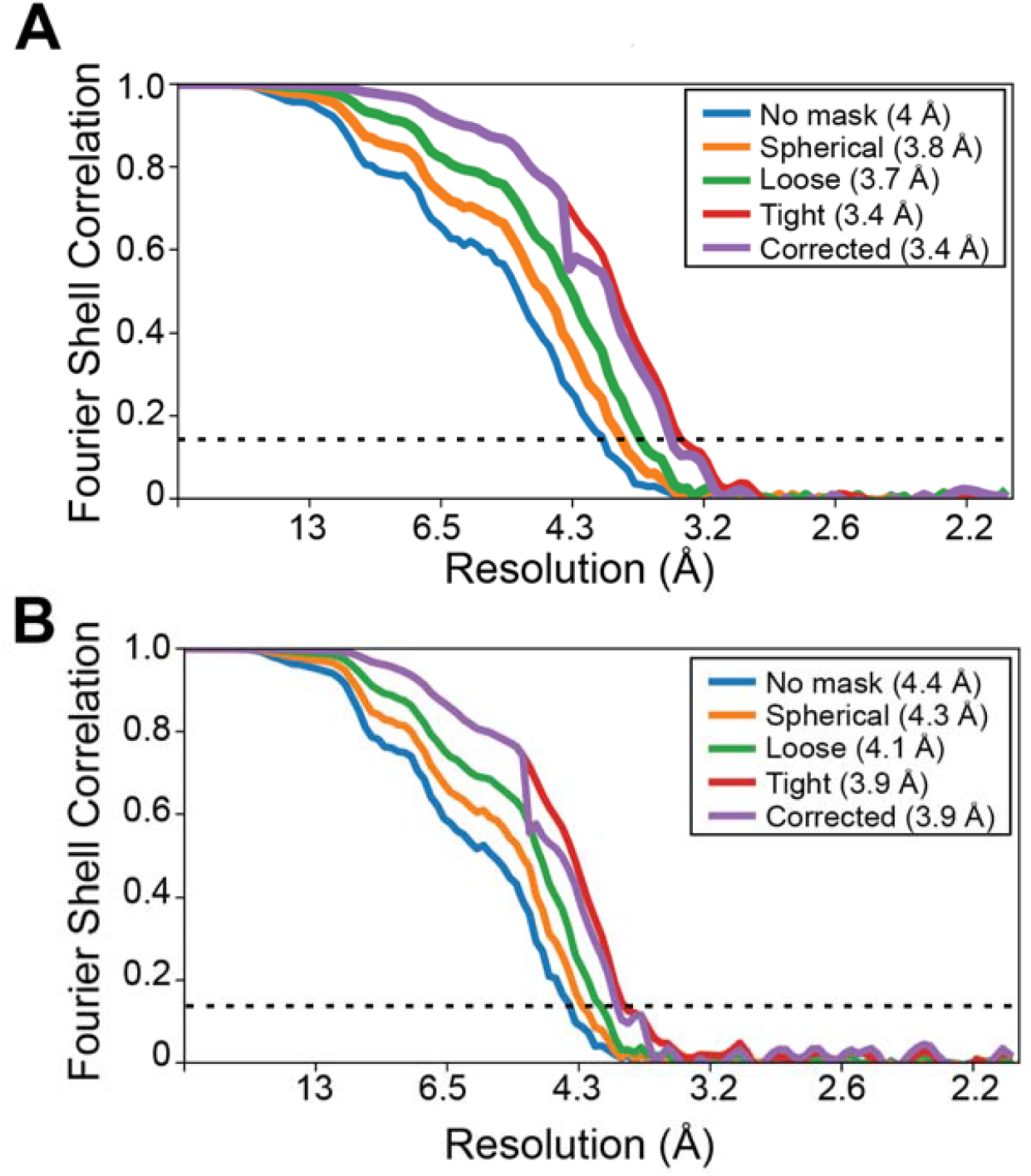
Calculation of map resolutions. Fourier Shell Correlation (FSC) curves for the refinement of the N (A) and M (B) states of TET, generated by cryoSPARC (*32*). The resolutions reported were estimated using half maps and gold standard FSC (GSFSC) of 0.143 and corrected using high-resolution noise substitution (*38*) to measure the amount of noise overfitting.

**Fig. S5.**
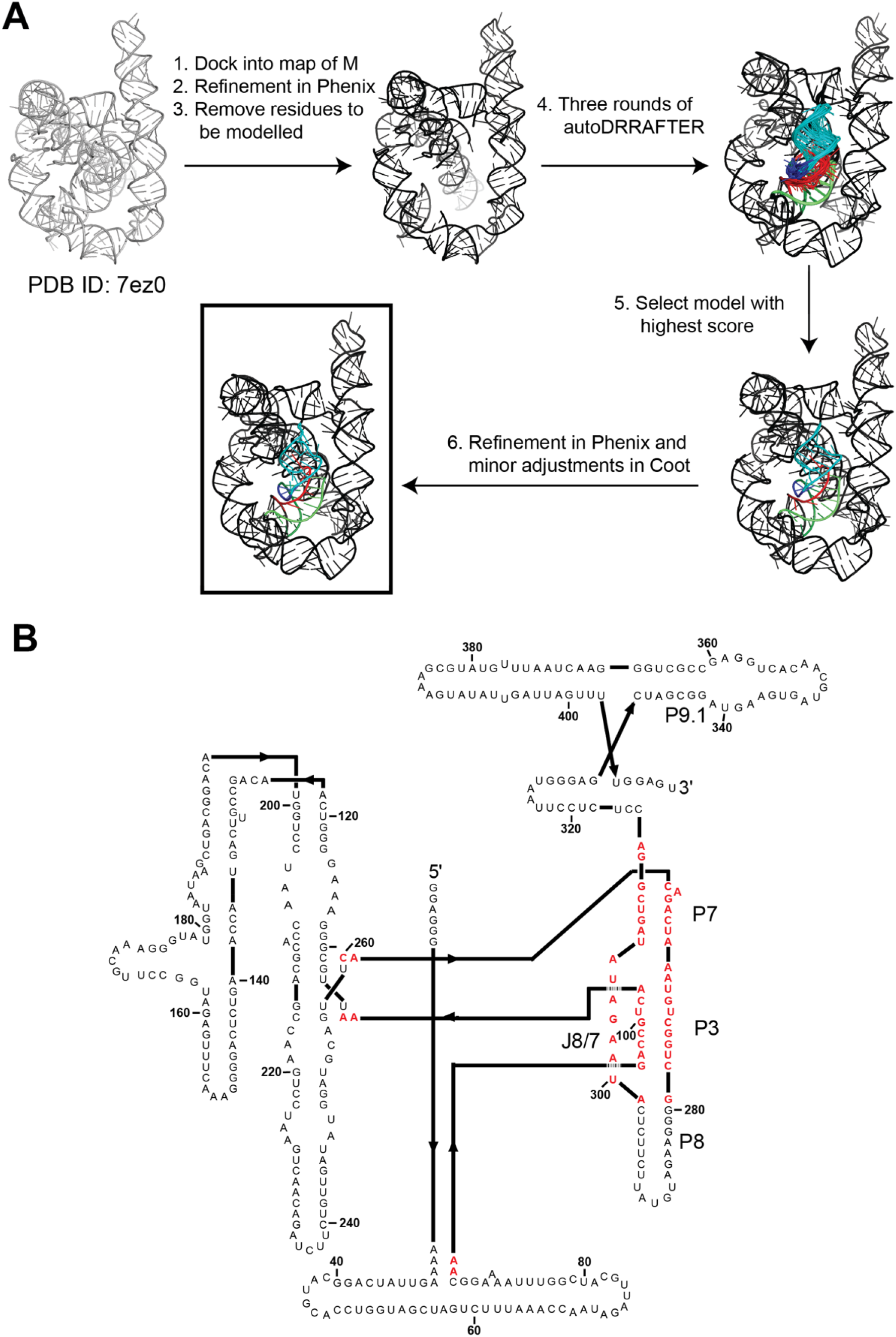
Structural modelling of M. (A) Workflow followed to model the structure of M using autoDRRAFTER (*13*), Phenix (*26*), and Coot (*27*). (B) Regions of TET modeled. Nucleotides that were removed from published structure of TET (PDB ID: 7ez0) and were modeled *di novo* by fragment assembly in autoDRRAFTER are colored red.

**Fig. S6.**
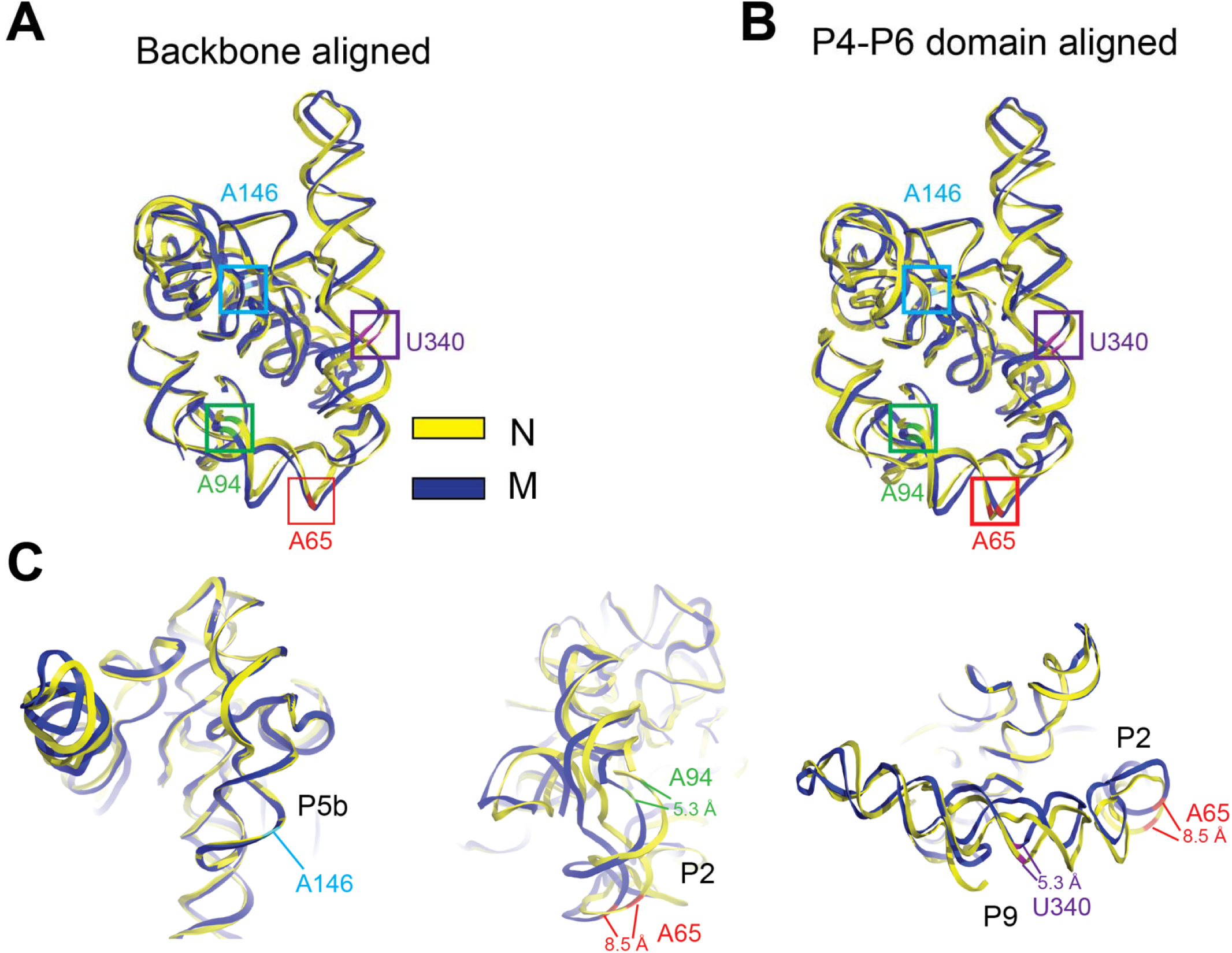
Differences in the relative position of peripheral domains between N and M. (A) Superposition of the backbones of N and M (RMSD_backbone_: 2.43 Å; 3758 atoms aligned). Regions that are distinct in the misfolded state were removed and did not contribute to the alignment. Nucleotides compared in (C) are colored and boxed. (B) Superposition of N and M based on the structural alignment of the P4-P6 domain (nts. 107 to 258; RMSD: 0.995Å). Colors are as in (A). (C) Superimposed structures (B) viewed from three angles. Distances measured between phosphates.

**Fig. S7.**
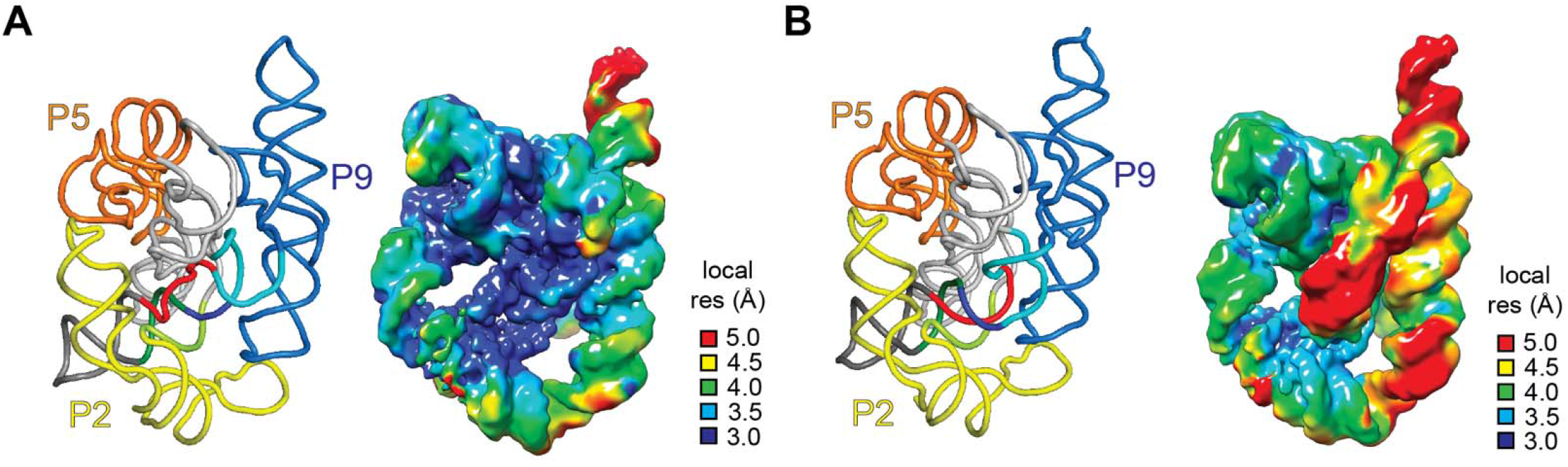
Local resolution of N and M cryo-EM maps. (A) Structure (PDB ID: 7ez0) and cryo-EM map of N colored according to its local resolution. (B) Structure and cryo-EM map of M colored according to its local resolution. Threshold for local Fourier Shell Correlation (FSC) resolution was set to 0.143.

**Table S1.** Data collection parameters and model statistics.

|  | TET M State | TET N State |
| --- | --- | --- |
| Microscope | Krios | Krios |
| Voltage (KeV) | 300 | 300 |
| Camera | Falcon 3 DED | Falcon 3 DED |
| Pixel size at detector (Å/pixel) | 0.5395 | 0.5395 |
| Total electron exposure (e-/Å<sup>2</sup>) | 32 | 32 |
| No. of frames collected during exposure | 46 | 46 |
| Defocus range (µm) | 0.8-2.0 | 0.8-2.0 |
| Automation software | SerialEM | SerialEM |
| Tilt angle (°) | 0 | 0 |
| Energy filter slit width (eV) | 20 | 20 |
| Micrographs collected (no.) | 6,222 | 6,222 |
| Total extracted particles (no.) | 4,272,705 | 4,272,702 |
| **Reconstruction** |  |  |
| Final refined particles (no.) | 98,071 | 92,828 |
| Point group | C1 | C1 |
| Resolution (Å) FSC: 0.143 | 3.9 | 3.4 |
| Map sharpening *B* factor (Å<sup>2</sup>) | 167.1 | 121.9 |
| Map sharpening methods | cryoSPARC<br>global sharpening | cryoSPARC<br>global sharpening |
| **Model Composition** |  |  |
| Protein | 0 | - |
| RNA | 386 | - |
| **Model Refinement** |  |  |
| Refinement package | Phenix | - |
| - Real or reciprocal space | Real | - |
| Model-Map scores |  |  |
| - CC (box) | 0.76 | - |
| - CC (mask) | 0.74 | - |
| - CC (volume) | 0.73 | - |
| - CC (peaks) | 0.66 | - |
| R.m.s deviations from ideal values |  |  |
| - Bond lengths (Å) | 0.001 | - |
| - Bond angles (°) | 0.412 | - |
| **Model Validation** |  |  |
| MolProbity score | 2.51 | - |
| Clashscore | 6.61 | - |

